# Estrogen receptor regulates hormone-induced growth arrest in a luminal A like breast cancer model

**DOI:** 10.1101/306662

**Authors:** Lacey Haddon, Sunny Hu, Hosna Jabbari, Brittney Loney, Zelda-Saidman Lichtensztejn, Richard Fahlman, Mary Hitt, Kirk McManus, Kelly Dabbs, John Mackey, Judith Hugh

## Abstract

Estrogen receptor positive (ER+) breast cancer has been divided into two subtypes, luminal A and luminal B, which differ in their ER expression and response to hormone therapy. The absence of luminal A cell lines means the extensive amount of *in vitro* work studying the response to hormones in ER+ breast cancers is biased for the luminal B subtype. We have developed a luminal A like cell model by increasing the ER expression in the MCF-7 cell line. Our results show that increased ER expression promotes an anti-proliferative response to estrogen through regulation of genes involved in the G1/S-phase transition of the cell cycle. Furthermore, increased ER expression increases ER-DNA binding in the absence of estrogen and regulates basal gene transcription by promoting DNA looping. These results provide novel evidence that the characteristic increased ER expression of luminal A tumors may promote a novel chromatin configuration that enables growth of these tumors in the absence of estrogen and enables gene repression in the presence of hormones.

---

Seventy-five percent of all invasive breast cancers express the estrogen receptor (ER) [1], a nuclear transcription factor that when activated by estrogen regulates cell growth [2]. ER+ tumors generally belong to one of two major intrinsic molecular subtypes, luminal A or luminal B [3], both of which are thought to be growth stimulated by estrogen and are treated with therapies that either target the ER or endogenous estrogen levels [4–5]. The likelihood of a clinical response to hormone therapy increases with increasing ER expression [6], thus patients with luminal A tumors that characteristically express high levels of ER have an excellent survival with hormonal manipulation alone [7]. Even with hormone therapy luminal B patients increasingly recur over time and benefit from chemotherapy [8–9]. The reproducible separation of these two intrinsic subtypes is one of the most important therapeutic challenges in breast cancer [10]. Even though there are several commercial tests available [11], the most accurate predictor of an excellent long-term outcome is a decrease in proliferation that occurs after a short exposure to hormonal therapy [12]. This suggests that the underlying hormone biology is key to predicting clinical outcome.

There is paradoxical clinical and laboratory evidence that estrogen is anti-proliferative [13]. Historically, estrogen was the treatment of choice for post-menopausal women with breast cancer [14] and in the landmark Women’s Health Initiative Study, estrogen only hormone replacement therapy was associated with a decreased incidence of breast cancer [15–16]. In the laboratory, after transfection of the estrogen receptor into either ER negative [17] or ER positive [18–19] cell lines, estrogen exposure results in growth suppression. We reasoned that the estrogen paradox was due to the two luminal subtypes having different responses to estrogen, with luminal A tumors being growth suppressed and luminal B tumors growth stimulated. Since all established ER+ cell lines subtype as luminal B [20–21], we developed a novel luminal A-like cell line to investigate the underlying mechanism(s) which differentiate the two luminal subtypes. We showed that increasing the expression of the ER, a luminal A characteristic, could change the transcriptional function of ER such that these tumors become growth suppressed by estrogen. We confirmed that increased ER expression alters ER-DNA binding patterns to more closely resemble hormone responsive tumors and further show that increased ER alters gene transcription via ER-mediated DNA reconfiguration.

## Results

### Increased ER expression leads to an anti-proliferative response to estrogen

We stably transduced a lentiviral plasmid with the ESR1 gene on a mEmerald doxycycline inducible promoter into the ER negative MDA-MB-231 (MDA-MB-231-ER) and ER positive MCF-7 (MCF7-ER) cell lines and determined the doxycycline concentration required to achieve a >20-fold increase in ER protein level (Figure 1a-b). Mock transfectants MDA-MB-231-EM and MCF7-EM were transduced with the empty mEmerald lentiviral plasmid and maintained their absent and low level of endogenous ER expression respectively after doxycycline induction (Figure 1a-b).

**Figure 1.**
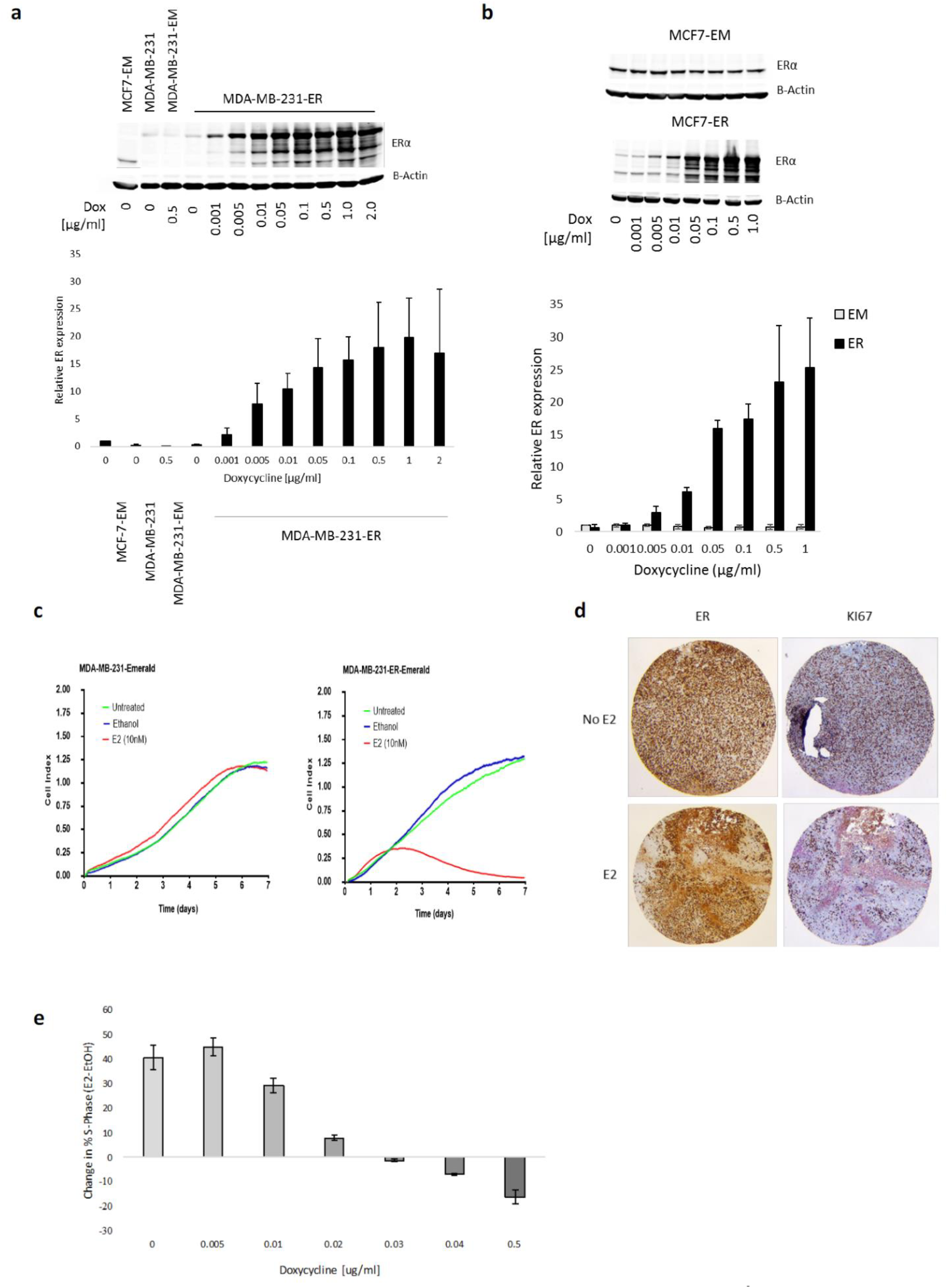
The level of ER expression induces a differential response to E2. (**a**) Representative western blot for ER protein expression in MDA-MB-231 parentals, MDA-MB-231-EM and MDA-MB-231-ER cells induced with varying doses of doxycycline. Relative ER expression was normalized to β-actin. (**b**) Representative western blot for ER protein expression in MCF7-EM and MCF7-ER cells induced with varying doses of doxycycline. Relative ER expression was normalized to β-actin. All experiments had n=3 and data is shown as mean ± s.d. (**c**) Growth (Cell index) of MDA-MB-231 cells transduced with (MDA-MB-231-ER-Emerald) or without (MDA-MB-231-Emerald) the ESR1 gene was measured in the absence of estrogen (untreated), vehicle control (ethanol) or in the presence of E2 (E2(10nM)) using Real-time cell analysis (RTCA). (**d**) Mouse xenografts derived from MDA-MB-231-ER-Emerald cells implanted in ovariectomized and immunocompromised mice treated with (E2) or without (no E2) subcutaneous estrogen capsules and stained for ER and Ki67 expression. (**e**) Change in % S-Phase in MCF7-ER cells induced with increasing concentrations of doxycycline and treated with or without 10nM E2 for 24 hours. Change in S-phase was calculated as the percentage of cells in S-phase for the E2 treated condition divided by vehicle control for each dose of doxycycline, n=3. Graph shows mean ± s.d. statistical significance was calculated by Student’s t-test. * P <0.05, ** P< 0.01 and *** P< 0.001.

To measure estrogen induced changes in cell growth, cells were adapted in estrogen-depleted media for 3 days, induced with doxycycline for 24 hours and then treated with 10nM estradiol (E2) for varying periods. After 5 days of E2-treatment the MDA-MB-231-ER cells showed a significantly decreased growth by real-time cell analysis (RTCA) with no effect on the mock transfected MDA-MB-231-EM (Figure 1c). Since MDA-MB-231 cells do not require estrogen supplementation to develop tumors in mice, we confirmed the anti-proliferative effect of estrogen using an *in vivo* model where MDA-MB-231-ER cells were xenografted into the mammary fat pads of immunocompromised, ovariectomized mice. After successful engraftment, doxycycline was added to the drinking water and two of four mice had estrogen capsules implanted subcutaneously. After 6 months, all four mice were sacrificed, and the tumors retrieved. The mice without the estrogen implant showed near universal expression of both ER and Ki67 (Figure 1d). Mice treated with estrogen had variable loss of ER throughout the tumor and Ki67 expression was only detected for cells in which ER expression was lost (Figure 1d). These results confirm that increased ER expression leads to a loss of cell growth in *in vitro* and *in vivo* models.

Because the MDA-MB-231 cell line derives from a basal-like intrinsic genetic background, we confirmed that the anti-proliferative effect of estrogen also occurs on a luminal background. MCF7-ER doxycycline titration experiments in the presence of estrogen showed there is a threshold of low ER expression that maintains the proliferative effect of estrogen and when the level of ER increases beyond this threshold the cells become anti-proliferative (Figure 1e). These results confirm the anti-proliferative effect seen in the MDA-MB-231-ER model is also present in MCF7-ER cells. More significantly, the ER-dose-dependent nature of this effect highlights a direct role for ER in the regulation of the anti-proliferative response.

### Direct ER-DNA binding regulates proliferative and anti-proliferative responses

ER is a ligand-activated transcription factor that regulates gene activation and repression by binding the DNA at a consensus palindromic estrogen response element (ERE) via two zinc fingers [22–23]. To determine whether the anti-proliferative effect is modulated by ER’s transcriptional function, we first inhibited transcription using Actinomycin D prior to E2 treatment. We found that Actinomycin D abrogated the proliferative (MCF-7-EM) and antiproliferative (MCF-7-ER) responses of these cell lines to E2 (Figure 2a). We then confirmed that this effect is mediated by ER-DNA binding by replacing the ESR1 gene in our lentiviral vector with a DNA binding mutant that contained three point-mutations (E203G, G204S and A207V) in the first zinc finger of the DNA binding domain [24]. We obtained stable transfectants (MCF-7-ERmDBD) with doxycycline induced ER expression similar to the levels seen in our MCF-7-ER cells (Figure 2b). Increased ER mutant expression in the MCF7-mDBD cells prevented the increase in basal proliferation seen in MCF7-ER cells in the absence of E2, the characteristic proliferative response to E2 as seen in the MCF-7-EM and the anti-proliferative effect of E2 seen in MCF7-ER cells (Figure 2c). This data suggests that ER regulates basal proliferation in the absence of E2, as well as the proliferative and anti-proliferative response to E2 via direct DNA binding and transcriptional activity.

**Figure 2.**
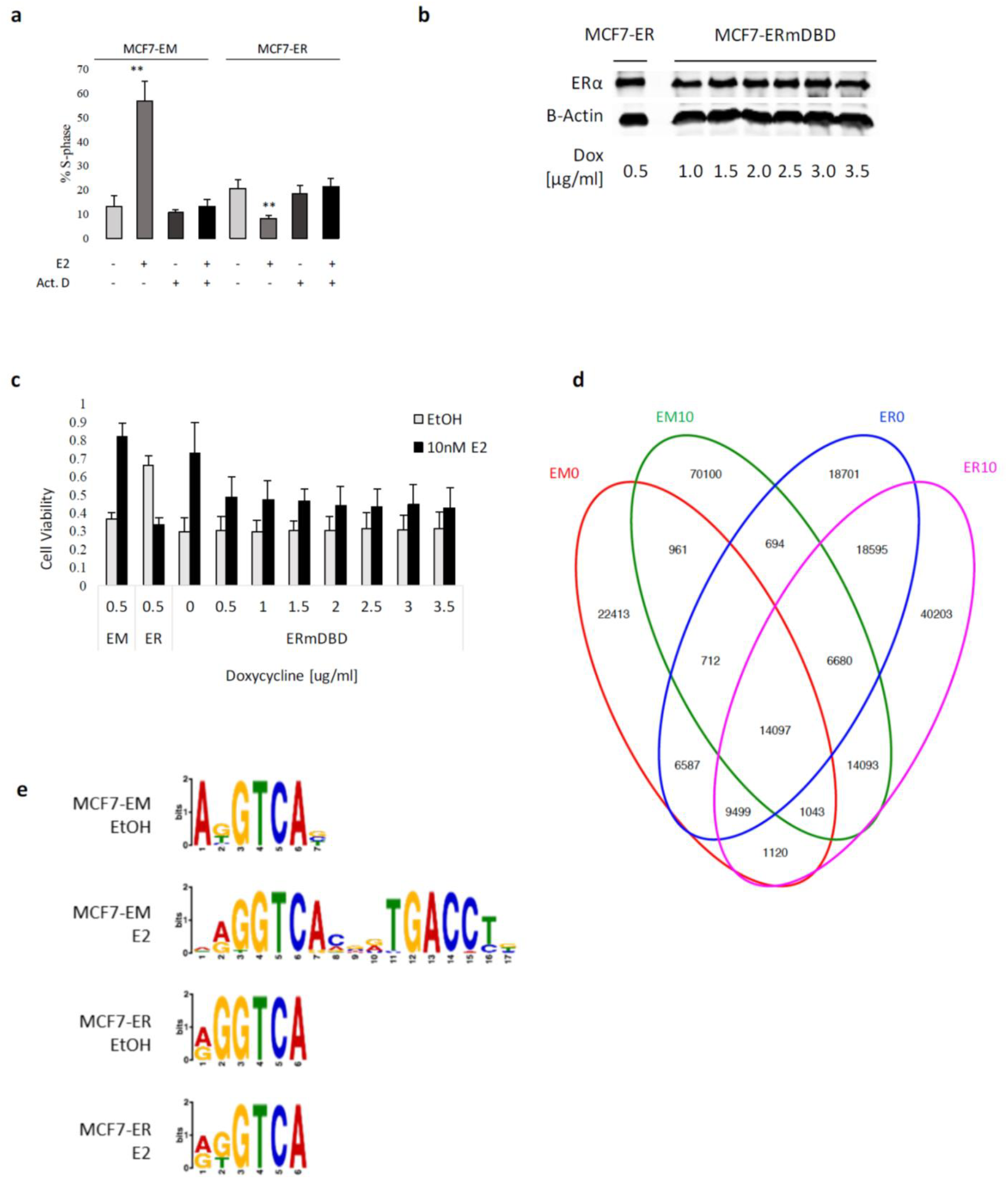
Increased ER expression promotes novel ER binding and transcription patterns in the presence of estrogen. (**a**) Percentage of MCF7-EM and MCF7-ER cells measured in the S-phase of the cell cycle by flow cytometry analysis in the presence of E2, actinomycin D or both. (**b**) Representative western blot for ER protein expression in MCF7-ER and MCF7-ERmDBD cells under varying doses of doxycycline induction. (**c**) Cell viability was measured in MCF7-EM (EM), MCF7-ER (ER) and MCF7-ER-mDBD (ERmDBD) cells treated with vehicle control (EtOH) or 10nM E2 under varying doses of doxycycline induction. All experiments had n=3. In a and c data is shown as mean ± s.d and analyzed by Student’s t-test. **P < 0.01. (**d**) Venn diagram of total ER peaks mapped by ChIP-Seq in MCF7-EM and MCF7-ER cells treated with vehicle control (EM0 and ER0) or 10nM E2 (EM10 and ER10), n=3. (**e**) Top motifs from MEME-ChIP analysis for MCF7-EM and MCF7-ER cells treated with vehicle control (EtOH) or 10nM E2 (E2).

To determine how increased ER mediates this differential response we investigated ER-DNA binding by chromatin immunoprecipitation followed by whole-genome sequencing (ChIP-Seq) for the MCF7-EM and MCF7-ER cell lines in the absence or presence of E2. We observed that increased ER expression leads to an overall increase in ER binding both in the absence and presence of E2 (Figure 2d). The most common motifs for the MCF7-EM cells in the absence and presence of E2 were a half ERE and the full ERE, respectively (Figure 2e). For the MCF7-ER cells, the most common motif was a half ERE, irrespective of the absence or presence of E2 (Figure 2e). The increase in ER binding both in the absence and presence of E2 suggests that ER may regulate the differential response to E2 through novel binding at genes involved in cell cycle regulation and proliferation in the MCF7-ER cells.

### Increased ER is not apoptotic but induces a G1 cell cycle arrest in the presence of estrogen

Previous investigations of the effect of estrogen on MCF-7 cells after long-term hormone depletion concluded the anti-proliferative effect of estrogen was due to the induction of apoptosis [25], however this estrogen-induced apoptotic response is not present in ER negative cells transfected with ER [26]. To determine if estrogen was inducing apoptosis we measured Poly (ADP-ribose) Polymerase (PARP) cleavage and the release of cytochrome C from the mitochondria using staurosporine as a positive control. We found no evidence of PARP cleavage or cytosolic cytochrome C after E2 treatment in either the MCF-7-ER or MCF-7-EM cell line (Figure 3d). These results support the clinical data showing that the treatment of ER+ tumors with tamoxifen, an anti-estrogen with documented estrogenic properties [27], induces growth suppression but does not increase or change the number of apoptotic events detected in these tumors [28].

**Figure 3.**
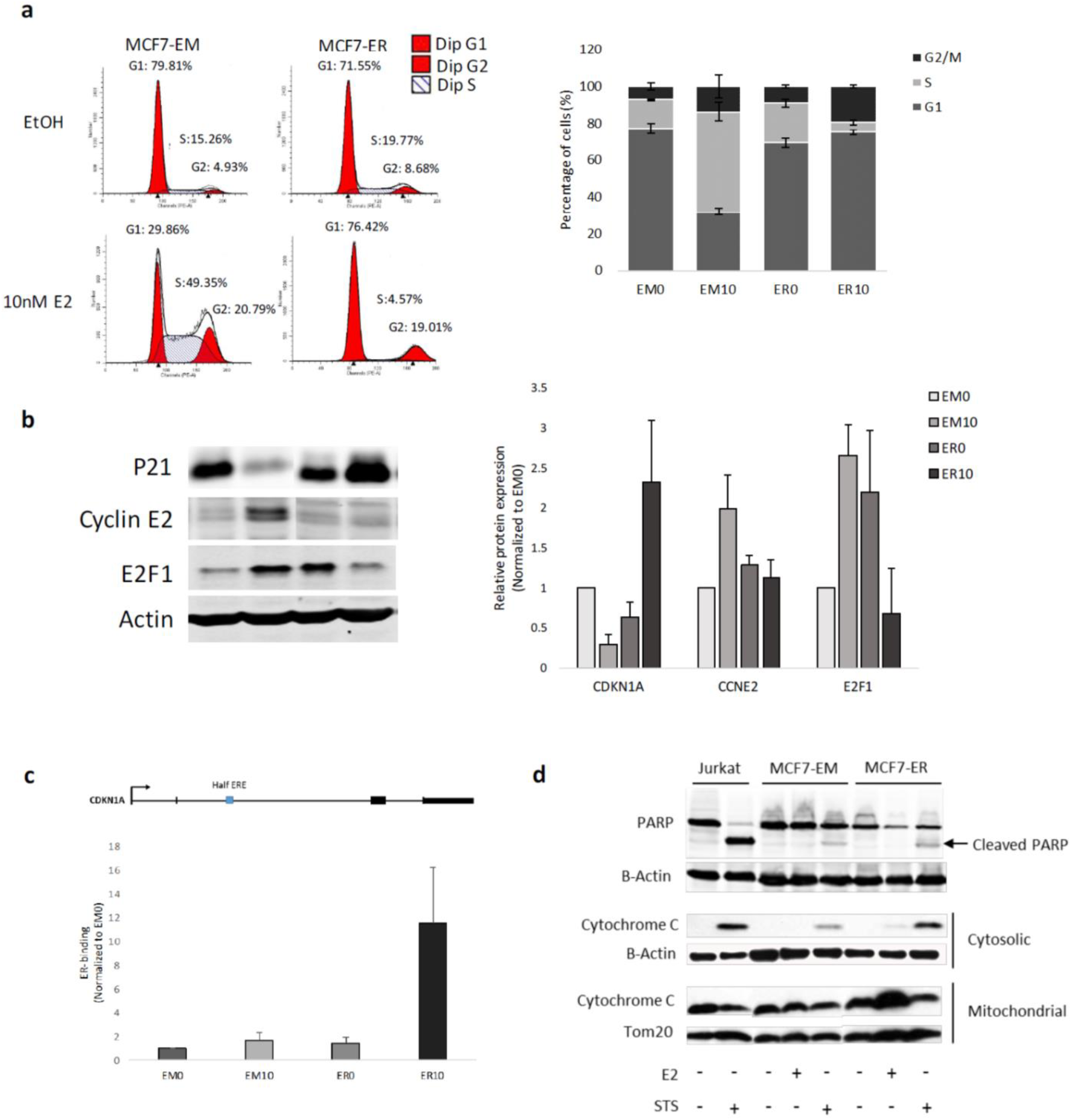
Increased ER expression causes cell cycle arrest in the presence of estrogen. (**a**) Cell cycle distribution of MCF7-EM and MCF7-ER cells treated with vehicle control (EtOH) or 10nM E2 for 24 hours was assessed by flow cytometry. Graph depicts the average percentage of cells in the G2/M, S and G1 phases. (**b**) Representative western blot for p21, cyclin E2, E2F1 and actin protein expression in MCF7-EM and MCF7-ER cells treated with vehicle control (EM0 and ER0) or 10nM E2 (EM10 and ER10). All samples were normalized to actin as loading control. Relative protein expression was calculated with EM0 set as 1. (**c**) ChIP-String validation of ER binding at an intragenic region of CDKN1A with samples normalized to EM0. (**d**) Representative western blot for PARP cleavage and cytochrome C protein expression in MCF7-EM and MCF7-ER cells were treated with E2 or staurosporine (STS) for 24 hours. β-actin and Tom20 served as loading controls. Jurkat cells were treated with STS as a positive marker for apoptosis. All experiments had n=3. In **a**, **b**, and **c** data is shown as mean ± s.d. In **b** the data was analyzed by Student’s t-test. * P <0.05.

Estrogen is known to promote the transition of MCF-7 cells from the G1 to S-phase of the cell cycle [29–30]. Consistent with previously published results [31] we found that E2 treatment relieves the G1 arrest of MCF7-EM after exposure to estrogen-free medium for three days (Figure 3a). Our results confirmed the work of Fowler et al. [32] in that MCF7-ER cells do not show a G1 arrest when kept in estrogen-free media but exhibited a basal increase in proliferation (Figure 3a).

When treated with E2 for 24 hours, MCF7-EM cells exhibit the characteristic proliferative response with a dramatic increase in S-phase (Figure 3a), whereas the MCF7-ER cells have a significant reduction in S-phase fraction (P= 0.003) and a significant increase in cells arrested in G1 phase of the cell cycle (P= 2.32^−05^) (Figure 3a). This G1 arrest was associated with a significant increase in the cyclin-dependent kinase inhibitor p21 which regulates the progression from G1 to S-phase through interactions with cyclin/cyclin dependent kinase (CDK) complexes (Figure 3b). ChIP-Seq analysis showed a novel ER peak within the intragenic region of p21 gene (CDKN1A), and ChIP-String analysis confirmed the presence of ER binding at this region only in the MCF7-ER cells in the presence of E2 (Figure 3c). The increase in p21 correlated to significant decreases in the transcription factor E2F1 and its target gene cyclin E, which mediates S-phase transition (Figure 3b). We did not detect ER-binding near the E2F1 gene in either of our cell lines with or without E2 (data not shown). These results suggest ER mediates the G1 arrest in MCF7-ER cells treated with E2 through direct transcriptional regulation of p21.

### The MCF7-ER cell line correlates with ER+ patients that respond to hormone therapy

To further validate our MCF7-ER cell line as a luminal A-like model we compared the peak sets from our MCF7-EM and MCF7-ER ChIP-Seq experiments against a previously published dataset of genome-wide ER-DNA binding profiles of ER+ breast cancer patients that were either responsive or non-responsive to tamoxifen. While a previous comparison of MCF-7 binding profiles correlated best (79.8%) with the tamoxifen non-responsive [33], the MCF-7-ER peaks correlated significantly with the with tamoxifen responsive breast cancer patients (Figure 4a) with the most common motif for these peaks identified as an ERE half site (Figure 4a). Longterm tamoxifen treatment can significantly increase the serum estrogen levels in breast cancer patients and tamoxifen itself has well-documented agonistic effects on ER [27, 34]. This estrogenic effect has been shown clinically through the increase in the progesterone receptor (PR) in patients treated with tamoxifen [35]. PR expression is up-regulated in the presence of estrogen through ER binding at the promoter region of the PGR gene [36]. Therefore, the binding profile seen in the tamoxifen responsive patients is likely an estrogenic response similar to that found in our MCF7-ER cell line. Preliminary results from our ongoing PRe-operative ESTradiOl Window of Opportunity Study in Post-Menopausal Women with Newly Diagnosed ER Positive Breast Cancer (PRESTO, NCT# 02238808) clinical trial have shown a luminal A patient who was treated with estrogen for 2 weeks prior to surgery had a decrease in ki67 on the surgical resection specimen (Figure 4b). These results support the use of our MCF7-ER cell line as an *in vitro* model that better represents the anti-proliferative response to hormones seen in luminal A tumors. We therefore utilized the MCF7-ER cell line to investigate the molecular function of ER in these tumors.

**Figure 4.**
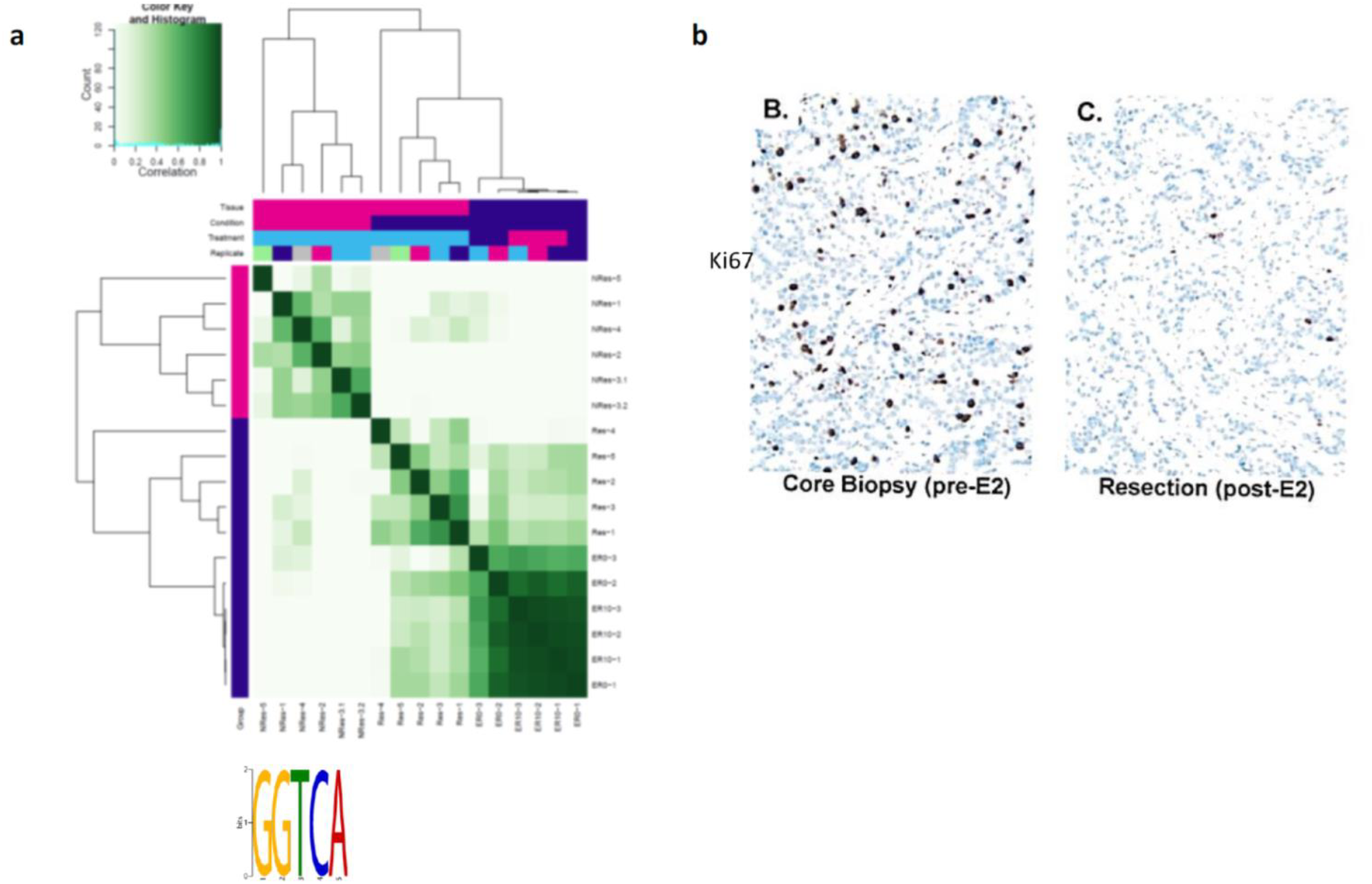
The MCF7-ER cell line correlates with ER+ patients that respond to hormones. (**a**) Correlation heat map for ER peaks obtained from MCF7-ER cells treated with vehicle control (ER0) or 10nM E2 (ER10) and tamoxifen responsive (Res) and tamoxifen non-responsive (NRes) patients from [33]. MEME-ChIP analysis of common peaks in MCF7-ER cells and tamoxifen responsive patients showed enrichment for the half-ERE (GGTCA). (**b**) Immunohistochemistry for Ki67 on core biopsy and resection samples after 2 weeks of estrogen treatment from a luminal A patient from the ongoing PRESTO clinical trial.

### Increased ER expression alters gene transcription via ER-mediated DNA reconfiguration

To assess the effects of increased ER expression on gene transcription, we measured transcriptome-wide changes in mRNA expression in the MCF7-EM and MCF7-ER cells with and without E2 by RNA-Seq. We searched for genes that were differentially regulated in the MCF-7-EM and MCF-7-ER cells using the MCF-7-EM in the absence of E2 as the baseline comparator. We found 72 genes that were (i) significantly increased with E2 in the MCF-7-EM (ii) increased without E2 in the MCF-7-ER and (iii) decreased with E2 in the MCF-7-ER (Supplemental Table 1) and confirmed this pattern of differential mRNA expression for five genes using qPCR (Figure 5a).

**Figure 5.**
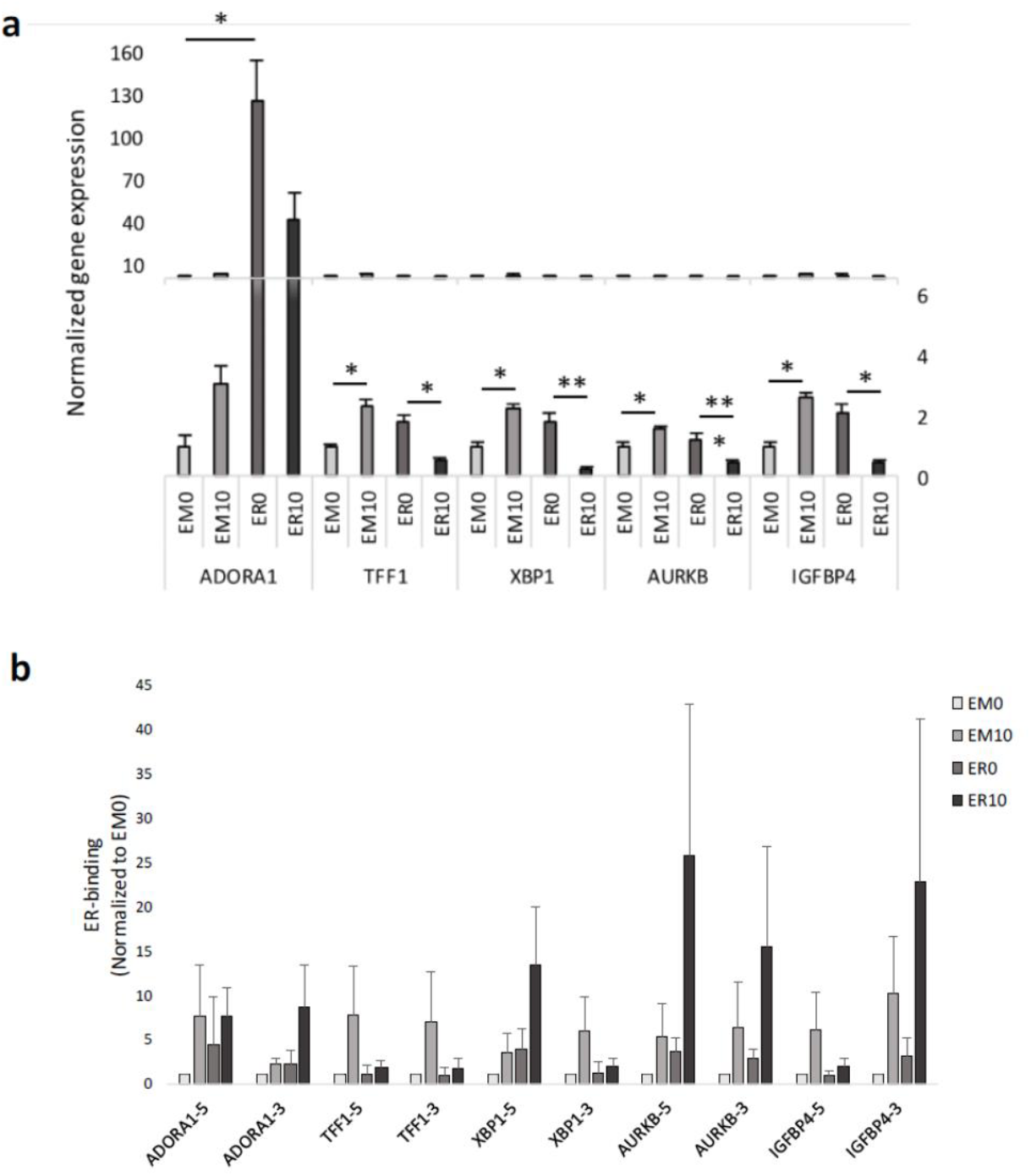
Changes in E2-mediated gene transcription involves direct ER-binding at known DNA loops. (**a**) Relative mRNA expression of the ADORA1, TFF1, XBP1, AURKB and IGFBP4 genes in MCF7-EM and MCF7-ER cells treated with vehicle control (EM0 and ER0) or 10nM E2 (EM10 and ER10) for 24 hours. mRNA expression was normalized to 3 housekeeping genes: PUM1, TBP, and RPL13A. (**b**) ER-binding at DNA regions associated with the 5’ and 3’ anchor regions (−5 and −3) of DNA loops associated with the ADORA1, TFF1, XBP1, AURKB and IGFBP4 genes in MCF7-EM and MCF7-ER cells treated with vehicle control (EM0 and ER0) or 10nM E2 (EM10 and ER10) was confirmed using ChIP-String analysis. All experiments had n=3. In a and b, data is shown as mean ± s.d. In a the data was analyzed by Student’s t-test. * P <0.05, ** P < 0.01 and *** P < 0.001.

ER binds to both proximal and distal enhancer regions [37] which can directly interact and promote gene activation via chromatin reconfiguration and the formation of large-scale DNA loops [38]. Using Homer [39] we confirmed that the MCF7-EM cells treated with E2 show enrichment of essential cofactors for ER-mediated chromatin reconfiguration: CTCF, FOXA1, GATA3 and AP-2γ [40] (Supplemental Table 2). Importantly, the MCF7-ER cells have a significantly greater enrichment of these motifs in the absence of E2 and maintain the enrichment of FOXA1, GATA3 and AP-2γ motifs after E2 treatment (Supplemental Table 2). The enrichment of these motifs in the MCF7-ER ChIP peaks supports the hypothesis that increased ER expression may promote the formation of DNA loops in the absence of estrogen.

We examined the 72 differentially regulated genes for evidence of looping by mapping our ChIP peak datasets against known ER-anchors obtained from the publically available ChIA-PET dataset for E2 treated MCF-7 (ENCODE). We chose 7 genes with ER peaks that overlapped with ChIA-PET interactions and validated the presence of ER-binding by ChIP-String analysis (Figure 5b). This confirmed that ER is bound to the mapped anchor regions in our MCF7-EM cells treated with E2; moreover, ER binding is maintained in MCF7-ER cells in the absence and presence E2 (Figure 5b).

We investigated the presence of a large-scale DNA loop (>2Mb) present in the TFF1 gene by fluorescence *in situ* hybridization (FISH) using a commercial fluorescently labelled BAC DNA probe set which corresponds to the 5’ and 3’ ER-anchor points of each loop (Figure 6a). Our results confirmed the presence of DNA looping in MCF7-EM cells treated with E2 (Figure 6b). Interestingly, DNA loops were increased in MCF7-ER compared to MCF7-EM cells in the absence of E2 (Figure 6b), and a similar amount of looping was maintained upon E2 treatment (Figure 6b). When taken together, the results from our MCF7-ER cell line provide the first evidence that increased ER expression promotes a response to estrogen that is mediated through the differential gene regulation via a novel configuration of ER-mediated DNA loops.

**Figure 6.**
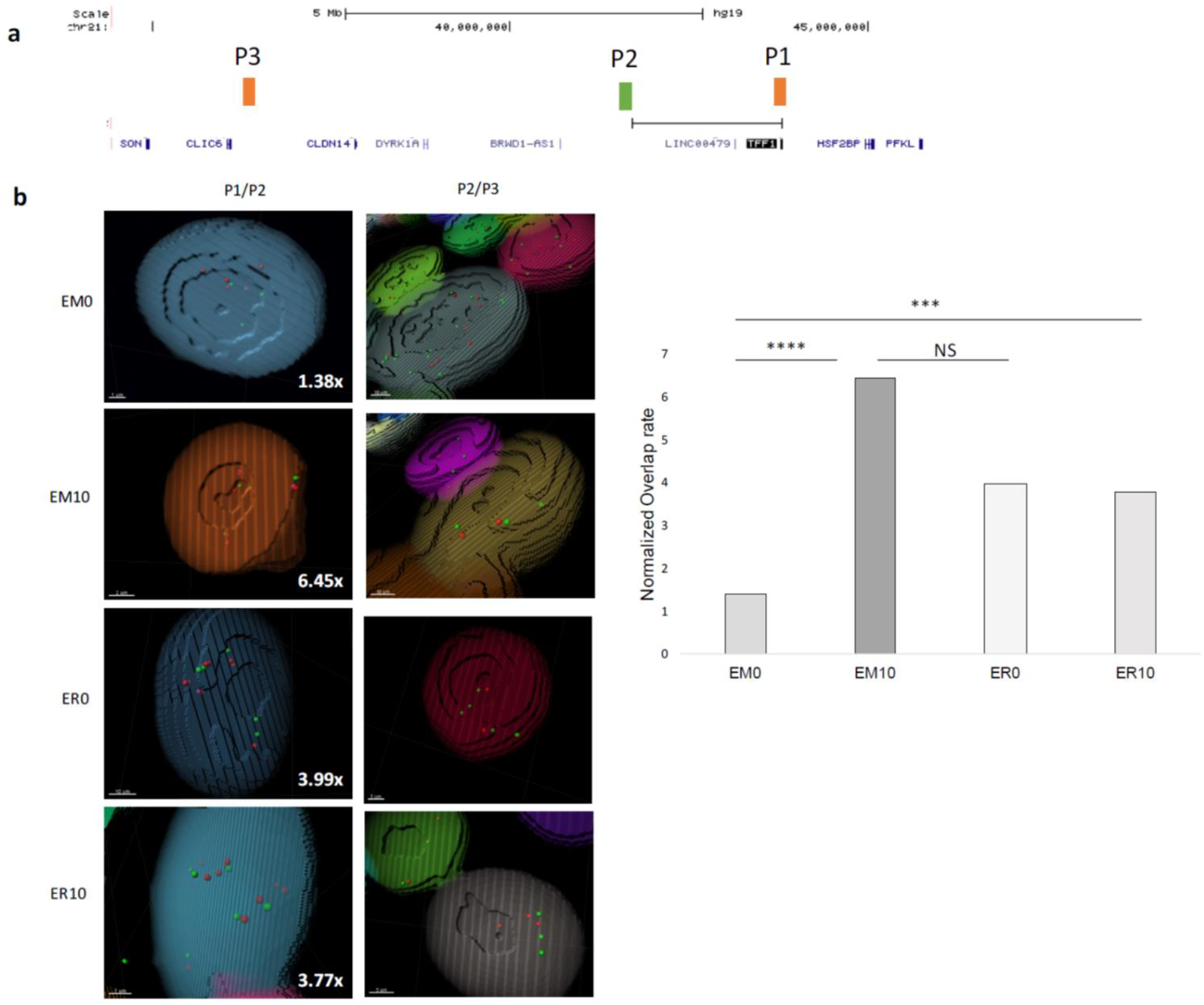
Increased ER expression influences the configuration of the TFF1 DNA loop. (**a**) P1 and P2 represent fluorescently labeled BAC probes near the anchor points associated with a >2Mb DNA loop near the TFF1 gene, and P3 is a control probe. (**b**) 3D confocal images from the IMARIS software showing BAC pairs are red and green spots and overlapping signals as red/pink. The probe overlap rate was normalized to the control pair (P2/P3). Data was analyzed by Fisher’s Exact test. *** P<0.001, **** P<0.0002, NS nonsignificant.

## Discussion

Molecular and clinical research has recognized at least two ER+ breast cancer subtypes with differing responses to hormones [41–42]. However, distinguishing the two subtypes has been problematic since all common ER+ cell lines genotypically represent the luminal B subtype and current molecular tests that rely heavily on the expression of proliferation genes are ~75% accurate at best [43]. Although an anti-proliferative response to E2 in cells that express high levels of ER is known [17–19], to the best of our knowledge we are the first to suggest that increased ER expression and the resultant E2-induced growth suppression can serve as a model for luminal A tumors. In this work, we used the ER-transfected MCF-7 cell line to explore the transcriptional impact of increased ER and compared our results against published clinical samples to further validate its use as an *in vitro* model for luminal A breast cancer. We also provide preliminary proof of a plausible transcriptional mechanism for this biological effect.

Luminal A tumors occur predominately in post-menopausal women [44], when serum E2 levels are nearly undetectable [45]. In our model, the ability of increased ER expression alone to alleviate the arrest of cells in G1 in the absence of E2 highlights a potential ER-mediated mechanism which enables luminal A tumors to develop in post-menopausal women. Our model and preliminary clinical studies showing an anti-proliferative effect of E2 in high-ER expressing cells would also explain the paradoxical anti-tumor results seen in the Women’s Health Initiative (WHI) and pilot studies of low-dose estrogen [46–47]. The absence of E2-induced apoptosis in this model is also consistent with published clinical findings [28] and our own window of opportunity clinical trial (results not shown). Finally, the similarities between the ER-DNA binding sites found by ChIP-Seq in the MCF7-ER expressing cells and the ChIP-Seq peaks of tamoxifen-responsive patients [33] are substantial evidence that a cell line expressing high levels of ER can serve as a relevant model for luminal A tumors.

Tamoxifen has been shown to prevent proliferation in MCF-7 cells through a G0/G1 arrest that is regulated by p21 and p27 [48]. Interestingly, this work and others have demonstrated that increased ER expression in the MCF7-ER cell line was enough to induce a p21-mediated G1 arrest in response to estrogen [19,26]. ER has been implicated in the transcriptional regulation of both the p21 (CDKN1A) and E2F1 genes through interactions with the transcription factors AP1 and/or Sp1 at the gene promoter regions [49–51]. Our ChIP-Seq experiments showed novel ER-binding at an intragenic region near the promoter of CDKN1A only in MCF7-ER cells treated with E2 but could not detect ER binding near the previously described motifs at the E2F1 promoter. These results suggest increased ER expression in our model enables binding of the CDKN1A gene and promotes the up-regulation of p21 in the presence of E2. Increased p21 expression would promote cell cycle arrest at the G1/S-phase check point through inhibition of the cyclin/CDK complexes which regulate the phosphorylation of retinoblastoma (Rb) protein which further releases E2F1 for transcriptional activity. This loss of E2F1 activity was shown through decreases in E2F1 protein, as well as the decrease in cyclin E, which is a target for E2F1 transcriptional activity and further promotes progression through the S-phase of the cell cycle [52]. These results are supported by previous work that found E2F1 was the major regulator of the differential response to E2 seen in MDA-MB-231 cells transfected to express high levels of ER [53]. Furthermore, the basal increase in proliferation seen in MCF7-ER cells in the absence of E2 is likely mediated through increased E2F1 activation as suggested by the increased levels of E2F1 protein in these cells. Previous research has shown the estrogen-independent growth that causes resistance to aromatase inhibitors in ER+ patients correlates with an E2F gene signature [54].

Our ChIP-Seq analysis found increased unliganded ER expression promotes novel ER-DNA binding and this was associated with a basal increase in gene expression. Furthermore, these peaks were enriched for motifs for transcription factors known to mediate DNA reconfiguration such as CTCF, FOXA1, GATA3 and AP-2γ. Comparison of the ER peaks present in the MCF7-ER cells without E2 against previously published ER-mediated loops in E2-treated MCF-7 cells suggested that increased ER expression may promote DNA looping in the absence of estrogen. As a proof-of-principle we investigated DNA looping at a >2Mb DNA loop mapped near the TFF1 gene using FISH. Our results provide novel evidence that increased ER expression in the absence of estrogen is enough to regulate chromatin reconfiguration. Interestingly, the TFF1 loop was maintained in the MCF7-ER cells treated with E2, however our gene expression data shows this gene becomes repressed under this condition. This suggests that increased ER expression may promote loops that mediate gene repression rather than activation in the presence of estrogen. Our ChIP-Seq data showed several novel ER binding peaks within the DNA loop region that are only present in the MCF7-ER cells in the presence of estrogen (data not shown). We hypothesize that these new E2 induced peaks may promote a novel loop configuration that causes gene repression rather that activation. Another ER-mediated loop at the TFF1 gene locus was shown to be maintained in MCF-7 cells treated with tamoxifen, which provides further support that loops can switch from an active to repressive state depending on the bound ligand [55]. The role of ER and the cofactors associated with repressive loops deserves further study.

Our preliminary results suggest that increased ER expression in the absence of estrogen can mediate changes in DNA binding and may promote and maintain DNA loops. Further experiments investigating the patterns of chromatin reconfiguration using the MCF7-ER model could reveal previously unmapped DNA loops and may provide novel insights into the biological mechanisms that regulate the response to hormones in luminal A tumors. This novel approach provides numerous opportunities for the future of ER+ breast cancer research, as it establishes an *in vitro* luminal A-like model which can be used to investigate differences in the ER+ subtypes which could led to the design of improved prognostic and diagnostic tools for use in the treatment and prevention of luminal A breast cancers.

## Methods

### Cell culture and estrogen treatment

MCF-7 parental cells (ATCC) were maintained in MEM (Gibco) containing 10% FBS (Gibco) and 10μg/ml bovine insulin (Sigma Aldrich). MDA-MB-231 cells were maintained in RPMI 1640 supplemented with 2 mM L-glutamine (Gibco) and 10% FBS. Stably transduced cell lines were maintained in media containing 10% Tet-free FBS (Clontech Laboratories, Inc), 500μg/ml geneticin (Gibco) and 1μg/ml puromycin (Sigma Aldrich). Prior to E2 treatment, cells were rinsed with PBS then adapted in phenol red-free media (Gibco) containing 10% carbon-stripped FBS (Sigma Aldrich) for three days. After adaptation, the MCF7-EM, MCF7-ER, MDA-MD-231-EM and MDA-MB-231-ER cell lines were induced with doxycycline for 24 hours, then treated with 10nM E2 or vehicle control (100% ethanol) for different time periods. The jurkat (Jneo) cells were maintained in RPMI 1640 supplemented with 10% FBS. The Lenti-X 293T cells were maintained in DMEM supplemented with 10% Tet-free FBS.

### ESR1 lentiviral constructs and stable cell line generation

ESR1 was amplified from ESR1 (NM 000125) human cDNA ORF clone (OriGene, Rockville, MD) by PCR using primers 5’-ATCCGCTAGCGCCACCATGACCATGACCCTCCACACCAAA-3’ and 5’-TCCGGAGGCTCGCGACCGTGGCAGGGAAACCCTCTGC-3’. The human ESR1 ORF was substituted for Dectin 1 into the Nhel and Sac1 sites of pmEmerald-Dectin1A-N-10 (kindly provided from Dr. Nicolas Touret, Department of Biochemistry, University of Alberta) to generate the pESR1-Emerald construct. The ESR1-Emerald was obtained by PCR from this plasmid using primers 5’-ATCCGCTAGCGCCACCATGACCATGACCCTCCACACCAAA-3’ (forward) and 5’-TCCGAGAATTCCGCTTACTTGTACAGCTCGTCCAT-3’ (reverse). In the forward primer, one Kozak consensus sequence was added to enhance expression. The PCR product was directly cloned into XbaI and EcoRI sites of the pLVX-Tight-Puro vector (Clontech Laboratories, Inc).

An ESR1 mutant containing three point mutations at E203G, G204S, A207V (ER-mDBD) was PCR amplified from pLVX-tet-on-tight-IRES-mcherry-ESR1-E203G, G204S, A207V-His (Biotechnology Creative Biogene) using the primers 5’-ATCCGGATCCGCCACCATGACCATGACCCTCCACACCAAA-3’ (forward) and 5’-CGGTGGATCCCCTCCGGAGCTCGCGACCGTGGCAGGGAAACCCTCT-3’(reverse), and cloned into the pLVX-Tight-Puro-ESR1-EM using the BamHI restriction sites. DNA sequence orientations and fidelities of the constructs were verified by both restriction enzyme digestion and full insert sequencing.

Lenti-X 293T cells were transfected with the LVX-Tet-on advanced (Clontech Laboratories, Inc), pLVX-Tight-Puro-EM, pLVX-Tight-Puro-ESR1-EM or pLVX-Tight-Puro-ESR1-mDBD-EM vectors using the Lenti-X HTX packaging system. Virus titers were collected for up to 48 hours. MCF-7 and MDA-MB-231 cells we co-transduced with the viral stocks supplemented with 6μg/ml polybrene according to the Lenti-X Tet-On Advanced Inducible Expression System User Manual (Clontech Laboratories, Inc).

### Western Blot Analysis and Quantification

Cells were lysed in RIPA buffer supplemented with fresh protease inhibitor cocktail and protein concentration was determined with Pierce BCA protein Assay Kit or Qubit 3.0. For mitochondrial fractionation experiments cells were lysed in digitonin lysis buffer (75mM NaCl, 1mM NaH_2_PO_4_, 8mM Na_2_HPO_4_, 250mM sucrose, 0.2mg/ml digitonin) and centrifuged at 14,000 rpm for 5 min. The supernatant was kept as a cytosolic fraction and the pellet was resuspended in triton X-100 lysis buffer (0.1% Triton X-100, 25 mM Tris PH 8.0) and kept as the mitochondrial fraction. For all experiments proteins were separated by SDS-PAGE and transferred to nitrocellulose membranes. Membranes were incubated with primary antibodies overnight at 4°C followed by 1-hour incubation with secondary antibody. Protein bands were scanned with Odyssey Infrared Imaging System (LI-COR Bioscience) and quantified using Image Studio (version 5.2).

### Antibodies

The antibodies specific for ERα (D8H8), PARP (46D11), cytochrome C (D18C7), p21Waf1/Cip1 (12D1) were obtained from Cell Signaling Technology. Antibodies against Cyclin B2 (R17985) were obtained from Abcam. Antibodies against β-actin (A5441) were obtained from Sigma Aldrich. The Tom20 antibody was a generous gift from Dr. Ing Swie Goping (Department of Biochemistry, University of Alberta). Alexa Fluor 700 goat anti-rabbit IgG (A21038) and Alexa Fluor 800 goat anti-mouse IgG (A32730) secondary antibodies were obtained from ThermoFisher Scientific.

### Real Time Cell Analysis

Real-time cell analyses (RTCA) (i.e. growth curves) were performed using an xCELLigence RTCA-DP (Acea Biosciences) as previously described [56]. Briefly, the RTCA-DP uses microelectrodes at the bottom of each well of an E-Plate 16 to measure increases or decreases in electrical impedance (termed Cell Index) that reflect increases or decreases in cell numbers, respectively. MDA-MB-231-Emerald and MDA-MB-231-ER-Emerald cells were treated with 10nM estradiol, ethanol (vehicle control), or left untreated (control), and 4000 cells from each condition were seeded into each well of an E-plate in quadruplicate and growth was monitored every 15min at 37°C. RTCA experiments were repeated at least once and graphs were generated by Prism V6 (GraphPad).

### Human Tumor Xenograft Studies

All animal experiments were approved by the Animal Care Committee at the Cross Cancer Institute (Edmonton, Alberta, Canada) and were carried out in accordance with guidelines of the Canadian Council on Animal Care. Contralateral xenografts were established by injecting ovariectomized female NIH-III mice (Charles River) in the abdominal mammary fat pads (MFPs) with 4 x 106 tumor cells in 50% Matrigel (MDA-MB-231-tER, right MFP; MDA-MB-231-tEM, left MFP). To induce expression of the tER and tEM coding sequences, mice were given doxycycline at 2 mg/ml in drinking water starting when tumors were palpable (day 32) and continuing for the duration of the experiment (day 73). One day after initiation of doxycycline treatment, 2 mice were implanted subcutaneously with beta-estradiol pellets (0.72 mg, 90-day time release, Innovation Research). All 4 mice were euthanized 11 weeks after tumor implantation. Tumors were excised, formalin-fixed and paraffin-embedded (FFPE) and tissue sections were sent to the Cross Cancer Institute (Edmonton, AB) for ER and Ki67 immunostaining. Blood was sampled by cardiac puncture at time of tumor harvest and serum estrogen levels were determined by electrochemiluminescence immunoassay (ECLIA) at the University of Alberta hospital.

### Flow Cytometry Analysis

Cells were trypsinized and centrifuged at 1400rpm. The pellets were washed and resuspended in PBS then fixed in 70% ethanol. Fixed cells were stained with propidium iodide (10μg/ml) for 30 minutes at room temperature. Cell cycle profiles were measured with BD LSRFortessa Special Order Research Product (SORP) (BD Biosciences) and analyzed using ModFit LT software.

### Chromatin Immunoprecipitation

ChIP was performed on MCF7-EM and MCF7-ER cells treated with vehicle control or 10nM E2 for 1 hour as previously described [57]. Briefly, treated cells crosslinked with 1% formaldehyde, sonicated, then incubated with ER antibody (HC-20, Santa Cruz-discontinued) overnight. ER-immunoprecipitates were collected with 1.5 mg protein A dynabeads, then reverse-crosslinked and digested with 40μg/ml proteinase K. DNA was purified with the ThermoFisher PCR purification Kit (K0702). A 1% input sample was collected for each ChIP experiment to serve as control. DNA was quantified using Qubit high sensitivity dsDNA reagents.

### ChIP Seq and Data Analysis

One ChIP-Seq replicate was performed by Active Motif on MCF-7-EM and MCF-7-ER cells treated vehicle control or 10nM E2 for 1 hour using their established ChIP protocol. Two additional ChIP sequencing libraries were prepared from the ChIP and Input DNAs using the NEBNext Ultra II DNA Library Prep Kit for Illumina (E7645S) and indexed using NEBNext Multiplex Oligos for Illumina (E7335S). DNA libraries were sequenced on Illumina’s NextSeq 500 using the NextSeq 500/550 High Output Kit (1x75 cycles) and reads were aligned to the human genome (hg19) using the Bowtie2 algorithm [58]. The aligned reads were filtered to retain only the uniquely mapped reads, which were then sorted, the duplicates were removed, and the remaining reads were indexed using SAMtools [59]. Peak locations were determined using the MACS2 algorithm (v1.4.2) with a cutoff p-value of 0.005 [60]. Published datasets for tamoxifen-responsive and resistant patients [33] were obtained from the Gene Expression Omnibus (GSE32222) and processed using the same bioinformatics protocol described above.

### ChIP-String Analysis

ChIP samples were obtained as described above. An nCounter custom ChIP-String codeset was designed by Nanostring Technologies against target sequences which correspond to significant peaks obtained in our ChIP-Seq experiments as well as regions devoid of ER binding to serve as negative controls (Supplemental table). ChIP and input DNA samples (1ng/μl) were denatured at 95°c for 5 minutes and then immediately cooled on ice. Hybridization buffer was added to the Reporter CodeSet and 8μl of this master mix was added to individual tubes. Ten microliters (total 10ng) of denatured DNA was added to the Reporter master mix, followed by 2μl of Capture ProbeSet (Nanostring Technologies). The samples were hybridized at 65°C overnight, then processed in the automatic Nanostring Prep Station. The fluorescent probes were counted the nCounter Digital Analyzer (Nanostring Technologies). All counts obtained from the nCounter were normalized to exogenous positive controls. The counts for the ChIP DNA sample were then normalized to the corresponding input samples for each probe set. To obtain the enrichment over background, each probe was normalized to the values obtained for the negative control regions. A final comparison was done to obtain the amount of ER binding relative to the baseline using the MCF7-EM cells in the absence of estrogen (set to 1). Three biological replicates of ChIP and Input DNA were used for ChIP-String analysis.

### RNA-Seq and Data analysis

Total RNA was extracted from MCF7-EM and MCF7-ER cells treated with vehicle control or 10nM E2 for 24 hours using the NucleoSpin RNA extraction kit, which included a DNase digestion step to remove potential contaminating DNA (Macherey-Negel). RNA quality was assessed using the Agilent Bioanalyzer 2100 and all samples had RIN scores from 8.6-10. cDNA libraries were made from 1μg total RNA using the NEBNext Poly(A) mRNA Magnetic Isolation Module (E7490) and NEBNext Ultra Directional RNA Library Prep Kit for Illumina (E7420) and indexed for sequencing using NEBNext Multiplex Oligos for Illumina. Libraries were sequenced on Illumina’s NextSeq 500 using the NextSeq 500/550 High Output Kit (2x75 cycles). The Fastq reads were aligned with UCSC genome sequence indexes and transcript annotation files for both hg19 and hg38 using the Tophat algorithm [61]. Differential gene expression analysis was done using Cuffdiff from the Cufflinks package [62].

### RNA quantitative RT-PCR

Total RNA was extracted as described for RNA-Seq. Reverse transcription followed by direct SYBR-green qPCR amplification was facilitated by the Power SYBR Green RNA-to-CT 1-Step Kit (Applied Biosystems). Taqman gene expression assays (Applied Biosystems) were purchased to measure the expression of genes of interest. Gene expression was normalized to three housekeeping genes: PUM1, TBP, and RPL13A.

### Comparison of ChIP-Seq peaks with known ER anchor regions

BED files from our ChIP-Seq analysis were uploaded to the University of California Santa Cruz (UCSC) genome browser along with a ChIA-PET dataset for MCF-7 cells immunoprecipitated with an ERalpha antibody generated by the ENCODE project (GSM970212). We used the combined datasets to locate ER peaks bound near each of the 73 differentially expressed genes from our RNA-Seq analysis. We selected an ER peak for further analysis if it met the following criteria: i) The peak was bound in the MCF7-EMs treated with 10nM E2 and the MCF7-ER cells in the absence of E2, ii) The ER peak overlapped with a mapped ChIA-PET anchor point near the gene in at least 2 of 3 ChIP replicates, and iii) The ChIA-PET interaction must span over 1Mb and maintain ER binding at both the 5’ and 3’ anchor points. The 1Mb threshold was set in order to enable detection of DNA loops using fluorescence *in situ* hybridization (FISH) as a validation method.

### Fluorescence *In Situ* Hybridization (FISH)

MCF7-EM and MCF7-ER cells were treated with or without 10nM E2 for 1 hour, trypsinized and resuspended in PBS. Cells were fixed in methanol and glacial acetic acid (3:1), dropped onto glass slides at 60°C then incubated in 2x SSC buffer at 37°C for 30 minutes. Slides were rinsed in dH_2_O and then dehydrated through 50%, 70%, 85% and 100% ethanol. Fluorescently labelled BAC probes (Empire Genomics) were mixed in hybridization buffer, added to slides and sealed with a coverslip. BAC probes were denatured at 73° for 5 minutes then hybridized at 37°C overnight. After hybridization, slides were washed in 0.4x SSC buffer (0.4x SSC and 0.3% IGEPAL, pH 7.0-7.5) at 73°C for 2 minutes, followed by 2x SSC wash buffer (2x SSC, 0.1% IGEPAL, pH 7.0 +/−0.2) at room temperature for 1 minute. Slides were rinsed in dH20 for 1 minute, then dehydrated through 70%; 85% and 100% ethanol. Slides were counterstained with DAPI I diluted 1:20 in VECTASHIELD mounting medium. Each condition was hybridized with the positive probe set against the 5’ and 3’ anchor regions of the TFF1 loop as well as a negative probe set which shared the 3’ anchor probe with the second probe located in a region with no looping at least >1Mb 3’ of the TFF1 loop (as described by [38]). Z-stack images were captured using the Zeiss LSM 710 laser scanning confocal with APD detectors and each image contained 200-300 nuclei. 3D Images were processed using IMARIS software and colocalized signals were counted using a threshold setting of 0.45 μM, which was shown to detect only the signals that were overlapping.

### Cell Viability Assay

MCF7-EM, MCF7-ER, and MCF7-ERmDBD cells were plated at 5x10^3^ in a 96-well plate and adapted in estrogen-free media as previously described. MCF7-EM and MCF7-ER cells were induced with 0.5μg/ml doxycycline and MCF7-ERmDBD cells were induced with increasing concentrations of doxycycline. After 24 hours, each cell line was treated in triplicate with 10nM E2 or ethanol control and incubated at 37°C and 5% CO2 for 5 days. Cell viability was assessed with TACS MTT assay (Trevigen) as per manufacturer’s protocol. Absorbance values were measured with the FLUOStar Omega microplate reader (BMG Labtech). Cell viability experiments were done in triplicate.

## Acknowledgements

We thank Dr. Kim Wood, Dr. Nicole Delaney, Debbie Tsuyuki, Kim Formenti, Dr. Russ Greiner for their work and support on this project. This work was made possible through the use of UCSC genome browser at http://genome.ucsc.edu/. This work was supported by Canadian Breast Cancer Foundation - Prairies/NWT Region, Lilian McCullough Endowed Chair in Breast Cancer Research, Canadian Cancer Society.

---

**Supplemental Table 1.** Increased ER expression leads to the differential expression of 73 genes in response to E2.

| Gene | EMO:EM10 |  | EMO:ERO |  | ERO:ER10 |  |
| --- | --- | --- | --- | --- | --- | --- |
|  | Log2 fold change | p-value | Log2 fold change | p-value | Log2 fold change | p-value |
| CD22 | 2.80 | 5.00E-05 | 4.42 | 5.00E-05 | -1.89 | 5.00E-05 |
| ITIH2 | 1.27 | 5.00E-05 | 1.82 | 5.00E-05 | -1.76 | 5.00E-05 |
| MYB | 2.61 | 5.00E-05 | 1.97 | 5.00E-05 | -2.75 | 5.00E-05 |
| RASGRP1 | 1.73 | 5.00E-05 | 2.65 | 5.00E-05 | -2.76 | 5.00E-05 |
| SLC17A9 | 2.68 | 5.00E-05 | 2.86 | 5.00E-05 | -2.97 | 5.00E-05 |
| SLC47A1 | 1.55 | 5.00E-05 | 3.19 | 5.00E-05 | -1.04 | 5.00E-05 |
| ADORA1 | 3.19 | 5.00E-05 | 5.14 | 5.00E-05 | -1.10 | 5.00E-05 |
| AMZ1 | 3.30 | 5.00E-05 | 3.45 | 5.00E-05 | -1.72 | 5.00E-05 |
| AURKB | 1.72 | 5.00E-05 | 1.04 | 5.00E-05 | -1.58 | 5.00E-05 |
| IGFBP4 | 2.34 | 5.00E-05 | 1.49 | 5.00E-05 | -2.18 | 5.00E-05 |
| OSGIN1 | 1.12 | 5.00E-05 | 2.60 | 5.00E-05 | -1.72 | 5.00E-05 |
| TFF1 | 3.11 | 5.00E-05 | 2.06 | 5.00E-05 | -1.01 | 5.00E-05 |
| XBP1 | 2.78 | 5.00E-05 | 2.89 | 5.00E-05 | -1.68 | 5.00E-05 |
| UGT2B15 | 5.09 | 5.00E-05 | 5.68 | 5.00E-05 | -2.66 | 5.00E-05 |
| RGS22 | 4.32 | 5.00E-05 | 3.69 | 5.00E-05 | -1.25 | 5.00E-05 |
| PDZK1 | 4.07 | 5.00E-05 | 4.02 | 5.00E-05 | -1.54 | 5.00E-05 |
| AGR3 | 3.47 | 5.00E-05 | 3.54 | 5.00E-05 | -3.70 | 5.00E-05 |
| LOC101928841 | 3.38 | 5.00E-05 | 3.23 | 5.00E-05 | -2.15 | 5.00E-05 |
| SYTL5 | 3.22 | 5.00E-05 | 3.29 | 5.00E-05 | -2.00 | 5.00E-05 |
| NPR1 | 2.69 | 5.00E-05 | 1.85 | 5.00E-05 | -2.37 | 5.00E-05 |
| RERG | 2.58 | 5.00E-05 | 1.77 | 5.00E-05 | -1.52 | 5.00E-05 |
| SCNN1B | 2.58 | 5.00E-05 | 2.04 | 5.00E-05 | -1.87 | 5.00E-05 |
| AREG | 2.46 | 5.00E-05 | 2.16 | 5.00E-05 | -2.07 | 5.00E-05 |
| SLC4A10 | 2.39 | 5.00E-05 | 1.22 | 5.00E-05 | -2.41 | 5.00E-05 |
| HS3ST3A1 | 2.32 | 5.00E-05 | 1.29 | 5.00E-05 | -1.81 | 5.00E-05 |
| SPOCK1 | 2.16 | 5.00E-05 | 2.11 | 5.00E-05 | -1.53 | 5.00E-05 |
| ASCL1 | 2.07 | 5.00E-05 | 1.23 | 5.00E-05 | -5.24 | 5.00E-05 |
| DSCC1 | 2.06 | 5.00E-05 | 1.08 | 5.00E-05 | -1.40 | 5.00E-05 |
| C1QTNF6 | 2.04 | 5.00E-05 | 1.16 | 5.00E-05 | -1.18 | 5.00E-05 |
| PRC1 | 1.45 | 5.00E-05 | 1.15 | 5.00E-05 | -1.16 | 5.00E-05 |
| PBK | 1.97 | 5.00E-05 | 1.00 | 5.00E-05 | -1.10 | 5.00E-05 |
| SPAG5 | 1.44 | 5.00E-05 | 1.05 | 5.00E-05 | -1.39 | 5.00E-05 |
| KRT16 | 1.97 | 5.00E-05 | 3.01 | 5.00E-05 | -1.28 | 5.00E-05 |
| RLN2 | 1.42 | 5.00E-05 | 1.70 | 5.00E-05 | -1.14 | 5.00E-05 |
| TMEM26 | 1.95 | 5.00E-05 | 2.76 | 5.00E-05 | -2.57 | 5.00E-05 |
| RNF223 | 1.41 | 5.00E-05 | 1.64 | 5.00E-05 | -2.30 | 5.00E-05 |
| SEMA3B | 1.36 | 5.00E-05 | 3.30 | 5.00E-05 | -1.86 | 5.00E-05 |
| MYBL2 | 1.79 | 5.00E-05 | 1.08 | 5.00E-05 | -1.62 | 5.00E-05 |
| PRR11 | 1.36 | 5.00E-05 | 1.24 | 5.00E-05 | -1.37 | 5.00E-05 |
| RNF183 | 1.75 | 5.00E-05 | 2.12 | 5.00E-05 | -1.43 | 5.00E-05 |
| SHCBP1 | 1.75 | 5.00E-05 | 1.16 | 5.00E-05 | -1.03 | 5.00E-05 |
| KRT6A | 1.32 | 5.00E-05 | 3.85 | 5.00E-05 | -1.20 | 5.00E-05 |
| HLA-DRA | 1.26 | 5.00E-05 | 2.22 | 5.00E-05 | -2.24 | 5.00E-05 |
| SKA1 | 1.68 | 5.00E-05 | 1.06 | 5.00E-05 | -1.00 | 5.00E-05 |
| DIRAS2 | 1.26 | 5.00E-05 | 1.14 | 5.00E-05 | -3.54 | 5.00E-05 |
| WDR62 | 1.67 | 5.00E-05 | 1.02 | 5.00E-05 | -1.02 | 5.00E-05 |
| UHRF1 | 1.25 | 5.00E-05 | 1.09 | 5.00E-05 | -1.46 | 5.00E-05 |
| SYTL4 | 1.67 | 5.00E-05 | 1.54 | 5.00E-05 | -1.24 | 5.00E-05 |
| MAD2L1 | 1.66 | 5.00E-05 | 1.03 | 5.00E-05 | -1.25 | 5.00E-05 |
| CCNB2 | 1.20 | 5.00E-05 | 1.06 | 5.00E-05 | -1.55 | 5.00E-05 |
| ORC1 | 1.64 | 5.00E-05 | 1.15 | 5.00E-05 | -1.65 | 5.00E-05 |
| HMMR | 1.20 | 5.00E-05 | 1.05 | 5.00E-05 | -1.00 | 5.00E-05 |
| CKAP2L | 1.60 | 5.00E-05 | 1.21 | 5.00E-05 | -1.25 | 5.00E-05 |
| ARHGAP11A | 1.50 | 5.00E-05 | 1.04 | 5.00E-05 | -1.21 | 5.00E-05 |
| ARHGAP11B | 1.18 | 5.00E-05 | 1.04 | 5.00E-05 | -1.28 | 5.00E-05 |
| NDC80 | 1.60 | 5.00E-05 | 1.09 | 5.00E-05 | -1.28 | 5.00E-05 |
| ST8SIA6 | 1.17 | 5.00E-05 | 1.09 | 5.00E-05 | -1.05 | 5.00E-05 |
| COL21A1 | 1.59 | 5.00E-05 | 1.87 | 5.00E-05 | -1.66 | 5.00E-05 |
| RAD51AP1 | 1.57 | 5.00E-05 | 1.17 | 5.00E-05 | -1.15 | 5.00E-05 |
| MMP13 | 1.12 | 0.00035 | 3.71 | 5.00E-05 | -1.80 | 5.00E-05 |
| OIP5 | 1.56 | 5.00E-05 | 1.12 | 5.00E-05 | -1.37 | 5.00E-05 |
| COL6A3 | 1.05 | 5.00E-05 | 1.57 | 5.00E-05 | -2.15 | 5.00E-05 |
| DTL | 1.49 | 5.00E-05 | 1.02 | 5.00E-05 | -1.66 | 5.00E-05 |
| KRT13 | 1.02 | 5.00E-05 | 2.38 | 5.00E-05 | -1.66 | 5.00E-05 |
| CENPU | 1.49 | 5.00E-05 | 1.09 | 5.00E-05 | -1.40 | 5.00E-05 |
| SHC4 | 2.04 | 0.0068 | 2.19 | 0.00355 | -2.43 | 0.03145 |
| CITED1 | 1.23 | 0.0018 | 3.73 | 5.00E-05 | -1.15 | 5.00E-05 |
| MSMB | 1.06 | 0.0009 | 1.10 | 0.00025 | -1.46 | 0.00025 |
| CDC25C | 1.03 | 0.0163 | 1.10 | 0.0088 | -1.34 | 0.0025 |
| VIM-AS1 | 2.09 | 5.00E-05 | 1.05 | 0.01395 | -2.40 | 0.0055 |
| LOC100506860 | 1.80 | 0.0004 | 1.86 | 0.0005 | -1.22 | 0.00295 |
| LINC00239 | 1.34 | 0.02095 | 2.31 | 0.0008 | -2.31 | 0.0018 |

**Supplemental Table 2.**
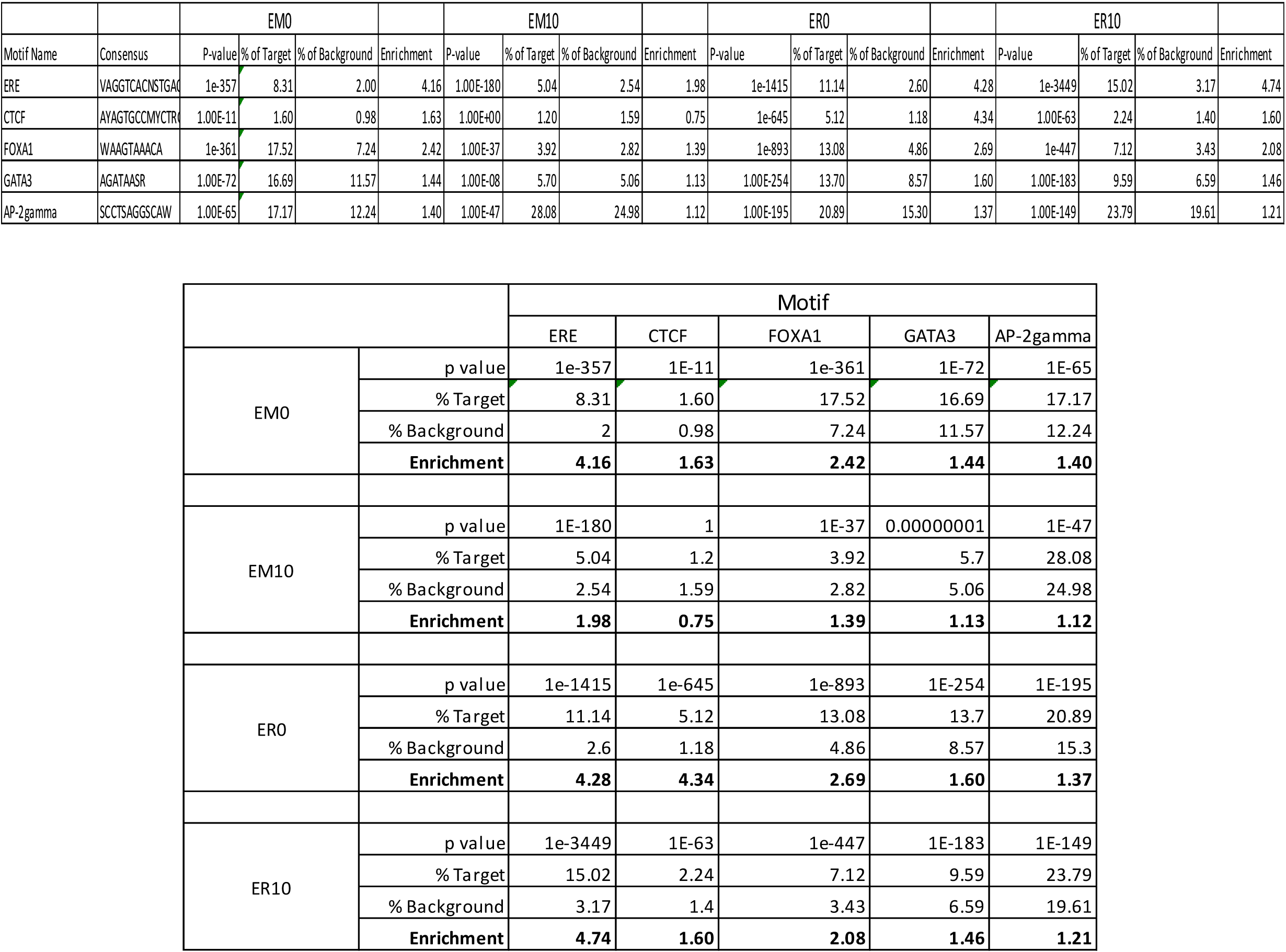
Motif enrichment by Homer analysis

